# Modulation of defence and iron homeostasis genes in rice roots by the diazotrophic endophyte: Herbaspirillum seropedicae

**DOI:** 10.1101/260380

**Authors:** L. C. C Brusamarello-Santos, D. Alberton, G. Valdameri, D. Camilios-Neto, R. Covre, K. Lopes, M. Z. Tadra-Sfeir, H. Faoro, R. A. Monteiro, A. B. Silva, W. J. Broughton, F. O. Pedrosa, R. Wassem, E. M. Souza

**Author notes:** **Corresponding author:** Address: Universidade Federal do Paraná/UFPR, Rua Coronel Francisco Heráclito dos Santos, s/n Caixa Postal 19046, 81531-980 – Curitiba. **E-mail of authors**: Brusamarello-Santos, L.C.C, Alberton, D., Valdameri, G., Camilios-Neto, D., Covre, R., Lopes, K., Tadra-Sfeir,1 M.Z., Faoro, H., Monteiro, R. A., Silva, A.B., Broughton, William J., Pedrosa, F.O., Wassem, R.

## Abstract

Rice is staple food of nearly half the world’s population. Rice yields must therefore increase to feed ever larger populations. By colonising rice and other plants, *Herbaspirillum* spp. stimulate plant growth and productivity. However the molecular factors involved are largely unknown. To further explore this interaction, the transcription profiles of Nipponbare rice roots inoculated with *Herbaspirillum seropedicae* were determined by RNA-seq. Mapping the 104 million reads against the *Oryza sativa* cv. Nipponbare genome produced 65 million unique mapped reads that represented 13,840 transcripts each with at least two-times coverage. About 7.4 % (1,019) genes were differentially regulated and of these 256 changed expression levels more than two times. Several of the modulated genes encoded proteins related to plant defence (e.g. a putative probenazole inducible protein), plant disease resistance as well as enzymes involved in flavonoid and isoprenoid synthesis. Genes related to the synthesis and efflux of phytosiderophores (PS) and transport of PS-iron complexes were also induced by the bacteria. These data suggest that the bacterium represses the rice defence system while concomitantly activating iron uptake. Transcripts of *H. seropedicae* were also detected amongst which genes involved in nitrogen fixation, cell motility and cell wall synthesis were the most expressed.

**Highlights:** RNASeq of *H. seropedicae* colonised rice roots showed remarkable regulation of defence, metal transport, stress and signalling genes. Fe-uptake genes were highly induced with implications in plant nutrition and immunity.

## Introduction

To answer the ever increasing demand for cereals, genetic improvement of rice and the concomitant development of bio-fertilisers are promising, low environmental-impact solutions. After water, the most limiting nutrient in plant development is nitrogen and nitrogenous fertilisers have been heavily used in rice cultivation (Ladha and Reddy, 2003). Heavy use of nitrogen fertilisers causes environmental damage including contamination of ground-water and the release of nitrogen oxides.

An alternative to the use of nitrogenous fertilisers is to employ plant-associated microorganisms that fix nitrogen. *Herbaspirillum seropedicae* is an endophytic diazotrophic that can colonise many plants and improve their productivity (reviewed by Monteiro *et al*., 2012; Chubatsu *et al*., 2012). Inoculation of rice with *H. seropedicae* increased root and shoot biomass by 38 to 54 % and 22 to 50 % respectively (Gyaneshwar *et al*., 2002) part of which was attributable to biological nitrogen fixation (James, 2000; Gyaneshwar *et al*., 2002; James *et al*, 2002; Roncato-Maccari *et al*., 2003). Pankievicz *et al.* (2015) showed that the nitrogen fixed by *H. seropedicae* and *Azospirillum brasilense* was rapidly incorporated into *Setaria viridis.* Other factors, including production of phyto-hormones by the bacteria stimulate plant growth and several authors have observed that the increase in biomass of inoculated plants is dependent on the plant genotype (Gyaneshwar *et al*., 2002; Sasaki *et al*., 2010).

Transcriptome based studies are powerful tools to detect differentially expressed genes and discover novel molecular processes (Nobuta *et al*., 2007; Mizuno *et al*, 2010; Xu *et al*, 2012, Xu-H *et al.* 2015; Wakasa *et al*, 2014; Magbanua *et al*, 2014; Shankar *et al*, 2016). Expression analyses (using EST sequencing and RT-qPCR) of rice roots inoculated with *>H*> *seropedicae* suggested that genes related to auxin and ethylene syntheses as well as defence are modulated by the microorganism in a cultivar dependent manner (Brusamarello-Santos *et al*., 2011). Here we used RNA-seq to profile the transcriptome of rice inoculated with H *seropedicae.*

## Materials and Methods

### Plant material and growth conditions

Testas were removed from seeds of rice *(Oryza sativa* ssp *japonica* cv. Nipponbare, kindly provided by the Instituto Riograndense do arroz, IRGA - Avenida Missões 342, Porto Alegre, RS, Brazil), then disinfected with 70 % (v/v) ethanol for 5 min followed by 30 min soaking in 8 % sodium hypochlorite (1 mL per seed) containing 0.1 % v/v Triton-X100. After rinsing 20 times with sterile water, the seeds were treated with 0.025 % (v/v) Vitavax-Thiram (Chentura, Avenida Nações Unidas 4777, Alto de Pinheiros, Sâo Paulo, SP, Brazil) fungicide solution and stirred (120 rpm) for 24 h in the dark at 30 °C. The seeds were then transferred to 0.7 % water-agar and left for two days to germinate after which the seedlings were inoculated with 1 mL of *Herbaspirillum seropedicae* strain SmR1 (10^8^ cells/seedling) for 30 minutes while control seedlings were treated with 1 mL of N-free NFbHP-malate medium (Klassen *et al*., 1997) (controls). Seedlings were washed with sterile water and transferred to glass tubes (25 cm long, 2.5 cm diameter) containing propylene beads and 25 mL of modified Hoagland’s solution (Hoagland and Arnon, 1950) without nitrogen (1mM KH_2_PO_4_, 1mM K_2_HPO_4_, 2mM MgSO4.7H2O, 2mM CaCl2.2H2O, 1mL/L micronutrient solution (H3BO3 2.86 g.L^−1^, MnCl2.4H2O 1.81 g.L^−1^, ZnSO4.7H2O 0.22 g.L^−1^, CuSO4.5H2O 0.08 g.L^−1^, Na2MoO4.2H2O 0.02 g.L^−1^) and 1mL.L^−1^ Fe-EDTA solution (Na2H2EDTA.2H2O 13.4 g.L^−1^ and FeCl3.6H2O 6 g.L^−1^)), pH 6.5-7.0. Plants were cultivated at 24 °C under 14 h light and 10 h dark for 3 days. *H. seropedicae* was cultivated in NFbHP malate medium containing 5 mM glutamate as the nitrogen source. Cells were shaken (120 rpm) overnight at 30 °C, then centrifuged, washed once with N-free NFbHP-malate and suspended in the same medium to OD_600_ = 1 (corresponding to 10^8^ cells.mL^−1^). Strain SmR1 (Souza *et al*., 2000) is a spontaneous streptomycin resistant mutant of strain Z78 (Baldani *et al*., 1986). Root colonisation was monitored by counting the number of colony forming units (CFU) per gram of fresh root. Approximately 100 mg of roots were washed in 70 % v/v ethanol for 1 min, 1% chloramine T for 1 min followed by three washes with sterile water (Dobereiner *et al*., 1995). The roots were then crushed with a mortar and pestle in 1 mL of NFbHP-malate and serial dilutions (10^−1^ to 10^−5^) were plated onto solid NFbHP-malate containing 20 mM NH4Cl. Two days later the cells were counted.

### RNA isolation and construction of libraries

The roots were separated from the aerial part and immediately stored in RNA later™ (Life Technologies, Foster City, CA, USA) Total RNA was extracted from roots of five rice plants using a RNAqueos kit (Ambion, Austin, TX, USA). Contaminating genomic DNA was eliminated with RNase-free DNase I (Ambion) for 30 min at 37 °C. Total RNA (7 to 10 μg) was depleted of ribosomal RNA by treatment with a RiboMinus™ Plant Kit for RNA-Seq (Invitrogen, Carlsbad, CA, USA). The integrity and quality of the total RNA was checked spectrophotometrically and by agarose gel electrophoresis. Whole Transcriptome Analysis RNA Kits™ (Life Technologies) were used on 500 ng purified RNA to construct the libraries and sequencing was performed in a SOLiD4 (Life Technologies) sequencer.

### Sequencing and analysis of short reads

SOLiD sequencing produced 161 million 50 bp reads that were analysed by SAET software (Applied Biosystems - Foster City, CA, EUA) to improve base calling, followed by quality trimming using the CLC Genomics Workbench (CLC bio, a QIAGEN Company, Silkeborgvej 2, DK-8000 Aarhus C, Denmark) (quality scores higher than 0.05 and reads with less than 20 bp were discarded). Then the reads were mapped to the rice genome database from the MSU Rice Genome Annotation Project (http://rice.plantbiology.msu.edu/) using the following parameters: a minimum length fraction of 95 %, minimum similarity of 90 % and only one hit. Differential expression was analysed using DESeq (Anders and Huber, 2010) from RobiNA software (Lohse *et al*., 2012). Genes covered at least twice were considered expressed and regulated when expression changed two times and the P-value was lower than 0.05.

### Quantification of mRNA levels using RT-qPCR

Reverse transcription quantitative PCR (RT-qPCR) analyses were used to evaluate gene expression under the conditions described above. Total RNA was isolated from rice roots using the TRI Reagent (Sigma, St. Louis, MO, USA) and contamination with genomic DNA was removed with DNase I (Life Technologies). The integrity and quality of the total RNA was confirmed by spectrophotometric analyses and electrophoresis. cDNA was produced from 1 μg DNase-treated total RNA using high-capacity cDNA reverse transcription kits (Life Technologies). The cDNA reaction was diluted 60 times before quantitative PCR using Power SYBR-Green PCR Master Mix on a Step One Plus Real Time-PCR System (both from Life Technologies). Primer sequences are listed in Table S1 and were designed with the Primer express 3.0 software (Applied Biosystems) and the NCBI primer designing tool using the genome sequence of *O. sativa* ssp. japonica cv. Nipponbare. Calibration curves for all primer sets were linear over four orders of magnitude (R^2^ = 0.98 to 0.99) and efficiencies were 90 % or higher. mRNA expression levels were normalised using the expression levels of actin 1, tubulin beta-2 chain (beta-2 tubulin) (Jain *et al*., 2006) and a hypothetical protein (protein kinase) (Narsai *et al*., 2010) using geNorm 3.4 software (Vandesompele *et al*., 2002). The relative expression level was calculated according to Pfaffl (2001). Three independent samples were analysed for each condition and each sample was assayed in triplicate.

## Results and Discussion

### Transcriptional analyses

*>H*> *seropedicae* enters rice roots via cracks at the points of lateral root emergence and later (three to 15 days) colonises the intercellular spaces, aerenchyma, cortical cells and vascular tissue (Elbeltagy *et al*., 2001; Gyaneshwar *et al*., 2002; James *et al*., 2002; Roncato-Maccari ei al., 2003). 14 days after inoculation we observed an increase of weight of roots and leaf but this increase was not statistically significant. The number of endophytic *>H*> *seropedicae* reached approximately 10^5^ to 10^6^ CFU per gram of fresh root weight one to two days after inoculation (DAI), with a peak at three DAI (data not shown). For this reason roots were collected for RNA-seq analyses three DAI when the population of *H. seropedicae* had stabilised in the intercellular spaces and xylem (Roncato-Maccari *et al*., 2003). Rice plants inoculated in parallel with the samples used for RNA extraction contained 4.2 × 10^5^ endophytic CFU.g^−1^ of fresh roots and 4.4 × 10^8^ epiphytic CFU.g^−1^ of root at three DAI.

Sixty-four percent of the reads (103,563,118) were finally used for mapping to the reference rice (www.rice.plantbiology) and *H. seropedicae* genomes. Illustration of RNA-seq analyses is shown in Figure 1 and the numbers of reads mapped to each reference genome are listed in Table 1. Mapping on the rice genome database (http://rice.plantbiology.msu.edu/) produced 22 million unique mapped reads representing 13,837 expressed transcripts.

**Figure 1.**
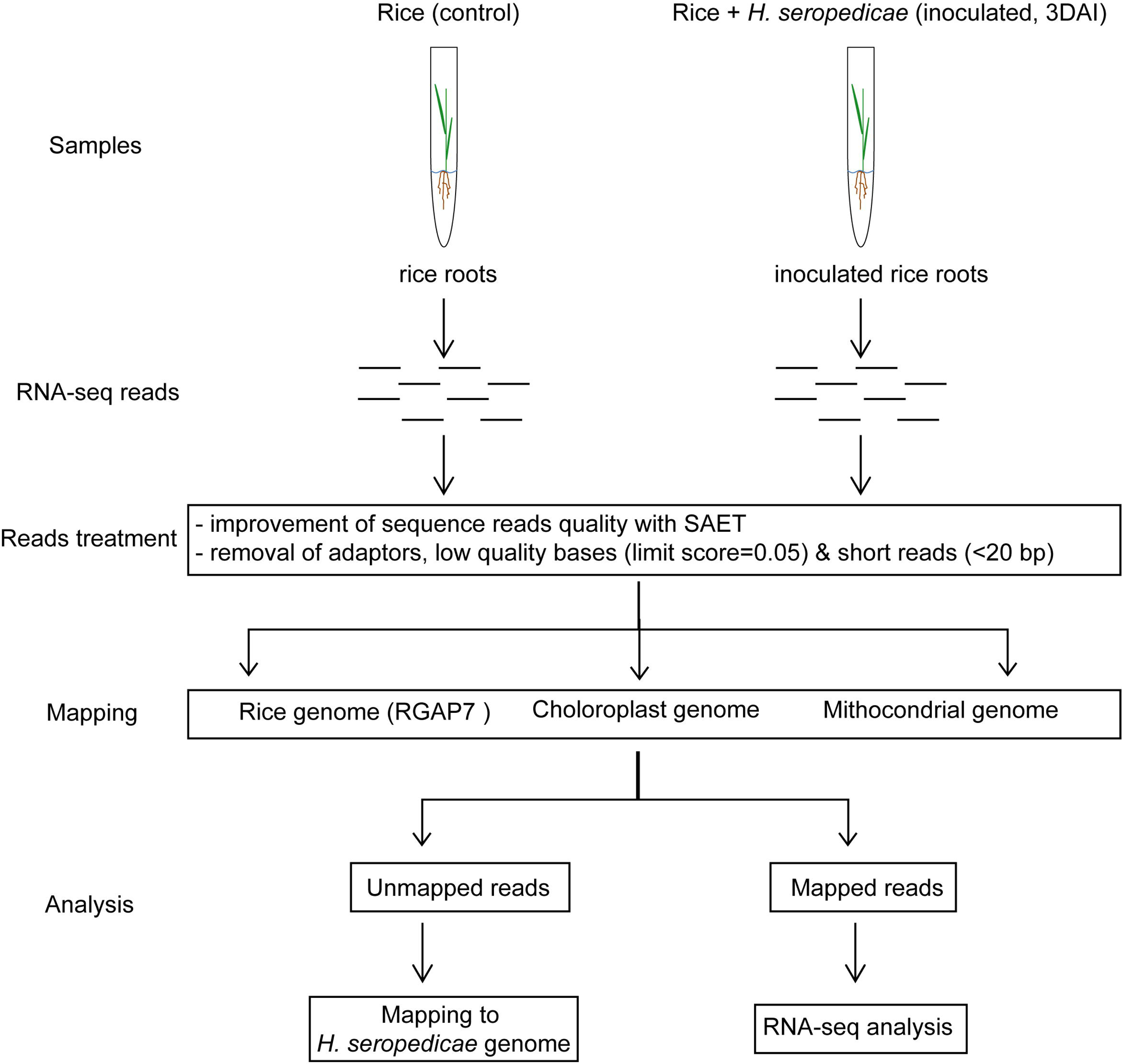
Diagram showing how rice roots were grown, inoculated with *H. seropedicae*, prepared for extraction of RNA and the RNA-seq analyses performed.

**Table 1.** Statistics of RNA-seq reads mapping

|  | **Control** | **Inoculated** |
| --- | --- | --- |
| **Total number of reads** | 101,587,721 | 59,591,404 |
| **Total number of reads after trimming** | 65,099,320 | 38,463,798 |
| **Reads uniquely mapped to RGAP7 (%)** | 12,255,809<br>(18.8) | 9,919,863 (25.8) |
| **Reads mapped uniquely to the rice**<br>**mitochondrial genome (%)** | 626,587 (0.96) | 253,771 (0.66) |
| **Reads mapped uniquely to the rice chloroplast**<br>**genome (%)** | 102,730 (0.16) | 36,915 (0.09) |
| **Total number of reads mapped to rice rRNA**<br>**(%)** | 41,744,369<br>(64.1) | 26,875,340 (69.9) |
| **Reads mapped to SmR1 genome (%)** |  | 16,878 (0.04) |
| **Total number of reads mapped (%)** | 54,803,125<br>(84.2) | 32,813,183 (85.3) |
| **Unmapped reads (%)** | 10,296,195<br>(15.8) | 5,650,615 (14.7) |

### Differentially expressed genes

Statistical analyses were performed using DESeq software (Anders and Huber, 2010) and comparison of non-inoculated with inoculated samples revealed 1,015 differentially expressed genes (P < 0.05). Amongst these, 255 had fold-changes higher than two and they were functionally categorised using MapMan (http://mapman.gabipd.org/) (Figure 2). Considering the number of regulated genes in relation to the number of expressed genes in each category, the main categories down-regulated by *H. seropedicae* were: metal handling (9.0%; 4/41), polyamine metabolism (9.1%; 1/11), secondary metabolism (9%; 15/167), hormone metabolism (4.7%; 11/233); stress (4.8%; 24/503) and nucleotide metabolism (4.9%; 6/123). Among genes up-regulated by bacteria were those involved in gluconeogenesis/glyoxylate cycles (28.6%; 01/07), polyamine metabolism (27.3%; 03/11), metal handling (24.4%; 10/41), fermentation (23.1%; 03/13) and nitrogen metabolism (18.8%; 03/16) (Figure 2). Some of these categories of regulated genes were analysed in more detail below.

**Figure 2.**
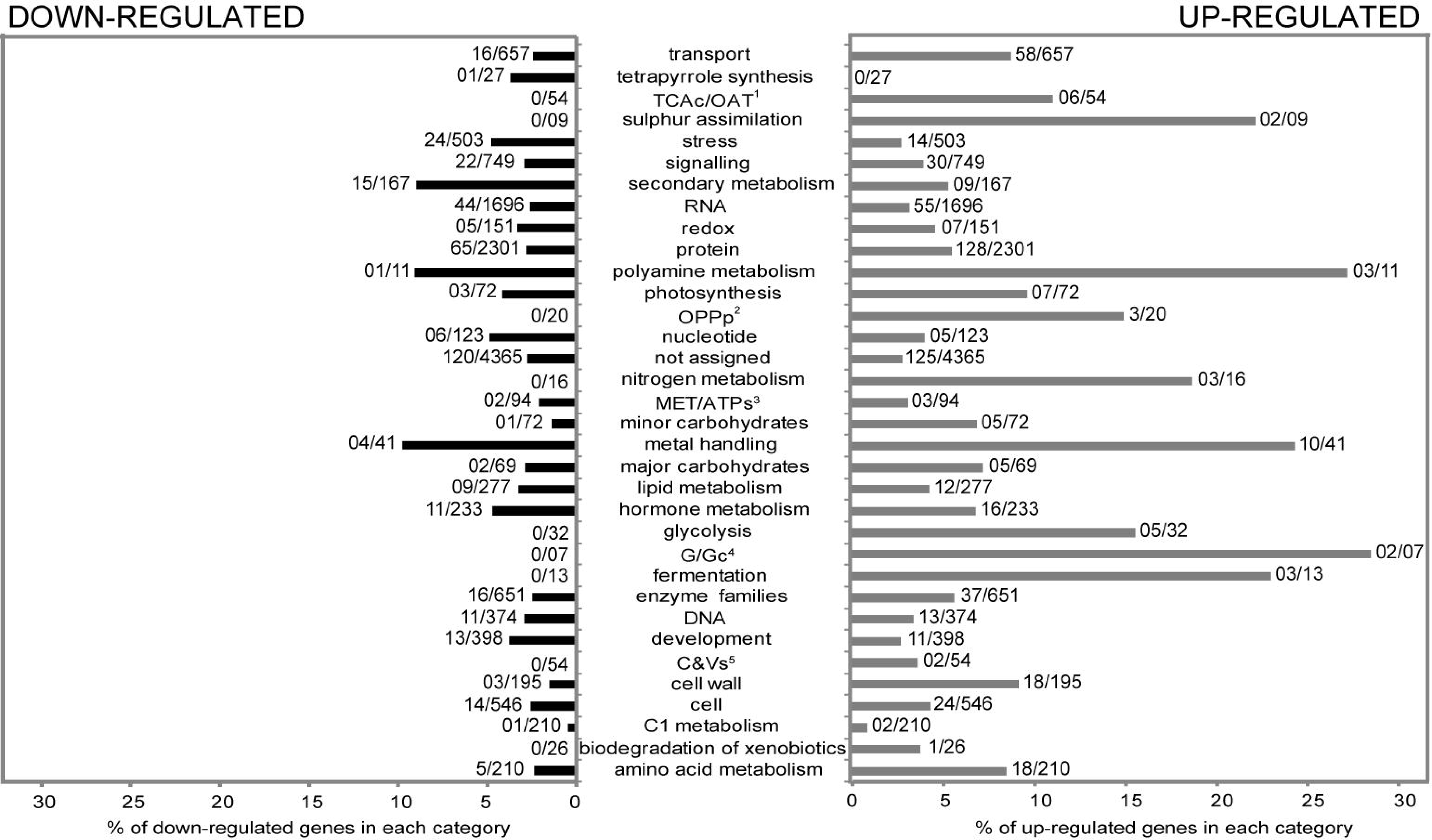
Transcripts differentially expressed in rice roots colonised by *H. seropedicae*. Up-regulated genes are shown in the right-hand column (in gray) with down-regulated genes to the left (in black). Numbers of regulated genes and total numbers of expressed genes are shown beside each column. Transcripts are grouped according to metabolic categories determined using MapMan.

### Biotic and abiotic stresses

Among the 255 differentially expressed genes (> 2fold), 59 were stress-related (30 repressed and 29 induced) in the following categories: secondary metabolites, hormone signalling, cell-wall, proteolysis, PR-proteins, signalling, transcription factors, redox-state, abiotic stress and peroxidases (Figure 3 and Table S2).

**Figure 3.**
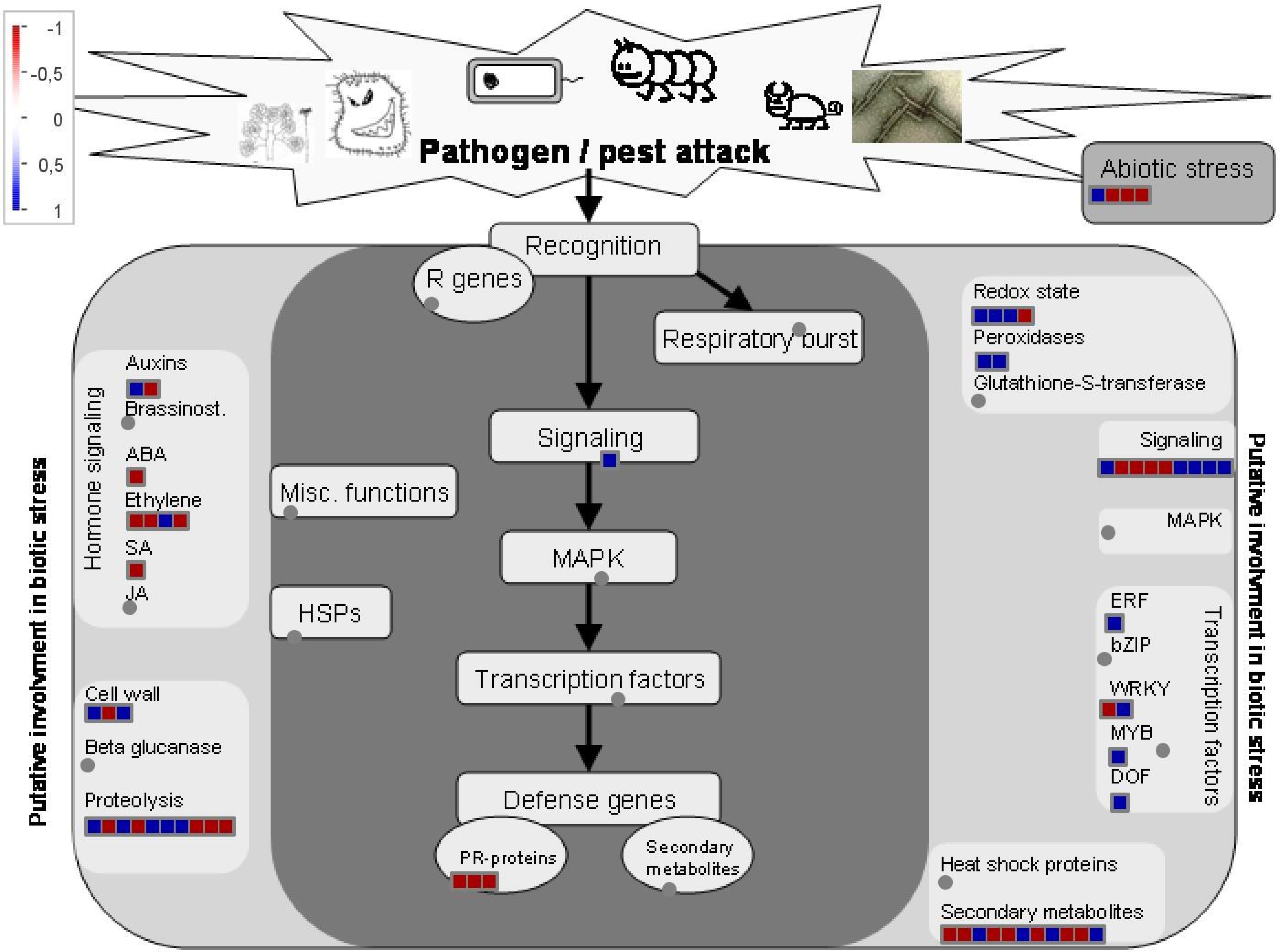
Biotic stress pathways regulated by *H. seropedicae* in rice roots. The scheme was constructed using MapMan (only genes with fold change ≥2, ≤2). Dark gray squares represent genes shown in these experiments to be involved in the biotic stress. On the left- and right-hand sides (shaded in light grey) are genes possibly involvement in biotic stress. Smaller squares represent up-regulated (blue) or down-regulated (red) genes.

#### Secondary metabolism

Amongst the secondary metabolic pathways with higher numbers of regulated genes were those involved in phenylpropanoid and isoprenoid synthesis (Table S2). Phenylpropanoids synthesised by deamination of L-phenylalanine are eventually converted to p-coumaric acid, a precursor of flavonoids and lignin (Ferrer *et al*., 2008; Hassan and Mathesius, 2012). *H. seropedicae* modulated expression of four genes involved in flavonoid synthesis. Amongst them, the gene encoding chalcone isomerase (CHI), which catalyses the synthesis of naringenin from tetrahydroxy-chalcone (Naoumkina *et al*., 2010), was repressed 2.1-fold. Naringenin is a key intermediate in the synthesis of other compounds including flavonol [flavonol synthase (FS), repressed 8.3-fold by *H. seropedicae*], and anthocyanins [dihydroxiflavonol 4-reductase (DRF), repressed 2.5-fold]. In addition, isoflavone reductase (IFR) gene, involved in isoflavone synthesis, was repressed 1.3-fold (P = 0.01). RT-qPCR of FS confirmed repression by *H. seropedicae* (Figure 4) but to a much lower extent (1.8-fold). Flavonols such as quercetin (Hassan and Mathesius, 2012) exhibit antimicrobial activity possibly by binding and inhibiting DNA gyrase (Plaper *et al*., 2003). Previously, Balsanelli *et al.* (2010) showed that narigenin has antimicrobial activity against *H. seropedicae*.

**Figure 4:**
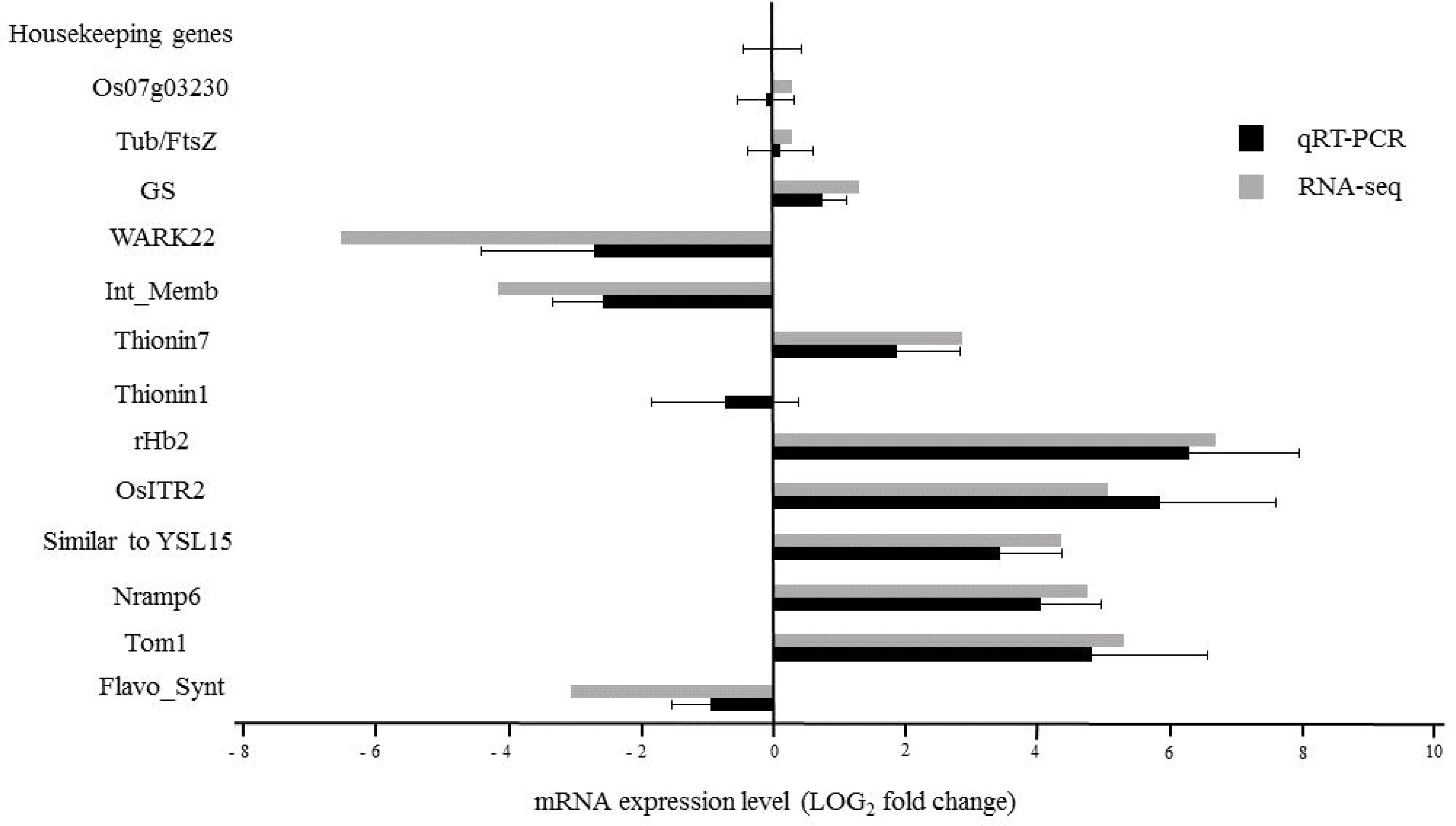
Confirmation of differential expression of rice genes by qRT-PCR and RNA-Seq. The results are average of three independent samples and error bars represent, the standard deviation. The reference genes used for the analysis were actin 1, tubulin beta-2 chain and conserved hypothetical protein.

Naoumkina *et al*. (2010) reported that flavonone-3-β-hydroxylase was induced in rice following infection with *Xanthomonas oryzae*. Nematode cysts stimulate isoflavone synthesis (including chalcone reductases, chalcone isomerase, isoflavon 2’-hydroxylase, isoflavones and isoflavone reductase synthase) in soybeans. Our data thus indicate that down-regulation of flavonoid/isoflavone synthesis by *H. seropedicae* is part of the attenuation of the defence system in rice roots that is necessary to host an endophyte.

As mentioned above *p*-coumaric acid is also a precursor of lignin which is assembled from monolignols (Vanholme *et al*., 2010). In this pathway cinnamoyl-CoA esters are converted into monolignols by two enzymes, cinnamoyl-CoA reductase (CCR) and cinnamyl alcohol dehydrogenase (CAD) (Boerjan *et al*., 2003). RNA-seq data showed that a gene (L0C_0s02g56700.1) encoding a cinnamoyl CoA reductase was induced 2.8-fold in rice inoculated with *H. seropedicae* (Table S2) while Kawasaki *et al*. (2006) reported that cinnamoyl-CoA reductase 1 (0sCCR1 - L0C_0s02g56460.1) is an effector of the small GTPase Rac that is involved in defence responses in rice (Kawasaki *et al*., 1999; Ono *et al*., 2001; Suharsono *et al*., 2002; Akamatsu *et al*., 2013). Transcriptome analyses of rice inoculated with the endophyte *Harpophora oryzae* or the pathogen *Magnaporthe oryzae* showed that a cinnamoyl CoA reductase (L0C_0s08g34280.1) like other enzymes in lignin synthesis was repressed in beneficial interactions and induced in pathogenic ones (Xu *et al*., 2015). L0C_0s08g34280.1 was also slightly repressed in our experiments (≈1.3 fold). It therefore seems as if a balance between induction and repression of defence-related genes allows *H. seropedicae* to survive inside rice tissues while concomittantly activating some defence responses to protect the plant from pathogens.

Isoprenoid (or terpenoid) synthesis was also modulated by *H. seropedicae* (Table S2 and Figure 5). Isoprenoids are derived from isomeric compounds including isopentenyl diphosphate (IPP) and dimethylallyl diphosphate (DMAPP) (Chen *et al*., 2011) via two pathways: the mevalonate pathway (MEV) and the methyl-erythritol phosphate pathway (MEP) (Figure 5). The MEV pathway occurs in the cytoplasm and the MEP pathway in plastids (Vranová *et al*, 2013) producing an estimated that 55,000 isoprenoid compounds. As a consequence, the biological function of these molecules is diverse and includes precursors of hormones, components of membranes, defence agents and carotenoids.

**Figure 5:**
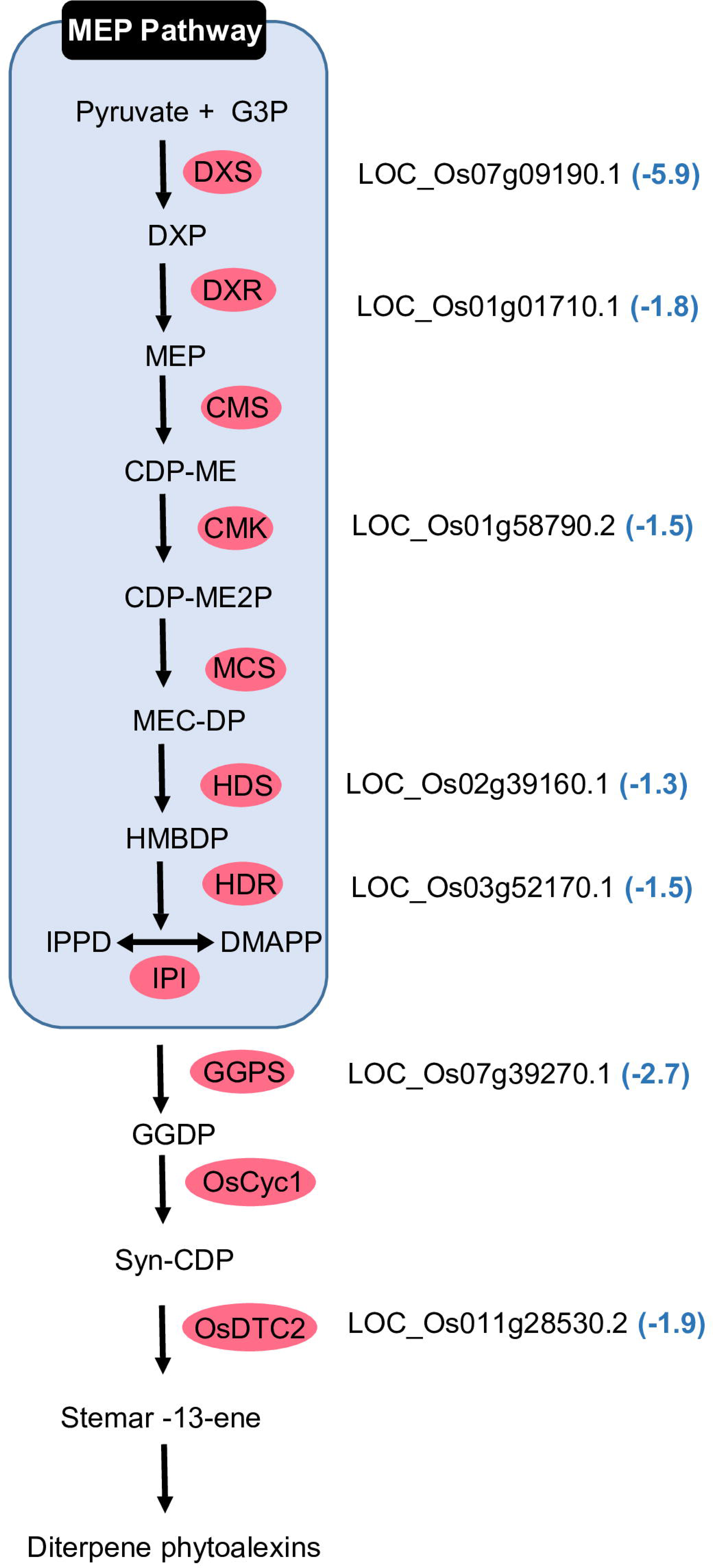
Isoprenoid synthesis genes down-regulated in rice roots colonised by *H. seropedicae*. The names of the genes differentially expressed are shown and the numbers in parentheses represent the fold change. The components of the MEP pathway leading to geranylgeranyl diphosphate synthesis and the diterpenoid-phytoalexin pathway are: G3P, glyceraldehyde-3-phosphate; DXP, 1-deoxy-D-xylulose 5-phosphate; MEP, 2-C-methyl-D-erythritol 4-phosphate; CDP-ME, 4-(cytidine 5-diphospho)-2-C-methyl-D-erythritol; CDP-ME2P, 2-phospho-4-(cytidine 5’-diphospho)-2-C-methyl-D-erythritol; MEC-DP, 2-C-methyl-D-erythritol 2,4-cyclodiphosphate; HMBDP, 1-hydroxy-2-methyl-2-butenyl 4-diphosphate; IPP, isopentenyl diphosphate; DMAPP, dimethylallyl diphosphate; GGDP, geranylgeranyl diphosphate; and CDP, copalyl diphosphate. Enzymes are indicated in rose coloured circles: DXS, 1-deoxy-D-xylulose 5-phosphate synthase; DXR, DXP reductoisomerase; CMS, CDP-ME synthase; CMK, CDP-ME kinase; MCS, MECDP synthase; HDS, HMBDP synthase; HDR, HMBDP reductase; IPI, IPP isomerase; GGPS, GGDP synthase; OsCyc1, syn-CDP synthase; OsCyc2, ent-CDP synthase; OsDTC2, stemar-13-ene synthase.

In rice, *H. seropedicae* repressed genes of the MEP pathway (Table S2 and Figure 5). The most repressed gene (5.9 times) encodes 1-deoxy-D-xylulose 5-phosphate synthase (DXS) the first enzyme of the pathway that synthesises 1-deoxy-D-xylulose 5-phosphate (DXP) from pyruvate and D-glyceraldehyde 3-phosphate. DXP is a precursor of antimicrobial compounds including phytoalexins. Chitin induces the MEP pathway in cultured rice cells and as a result phytoalexins accumulate (Okada *et al*, 2007). Furthermore treatment of the cells with inhibitors of the enzymes DXS and 1-deoxy-D-xylulose 5-phosphate reductoisomerase (DXR) impeded chitin-dependent accumulation of phytoalexin. The authors suggest that activation of the MEP is required to meet the demand for isoprenoid and phytoalexin synthesis in infected cells. In our experiments and those of Drogue *et al*. (2014), the gene encoding stemar-13-ene synthase (0sDTC2), an enzyme involved in synthesis of phytoalexins in rice, was repressed 1.9-fold in *H. seropedicae* treated rice roots (Figure 5) and twofold when infected with *Azospirillum* spp. (Drogue *et al*, 2014). It thus seems as if *H. seropedicae* represses production of defence-related isoprenoids in rice roots perhaps to allow the bacteria to enter and rapidly colonise the intercellular spaces and xylem.

#### Oxidative stress responses

In this study *H. seropedicae* modulated expression of five redox state genes including those for non-symbiotic hemoglobin 2 and peroxiredoxin. 0ne peroxiredoxin gene (L0C_0s07g44440.1) induced 3.7 and the other L0C_0s01g16152.1 was repressed 2.1 fold. Peroxiredoxins are a group of H202-decomposing antioxidant enzymes related to the redox state. In addition to the reduction of H_2_0_2_, peroxiredoxin proteins also detoxify alkyl hydroperoxides and peroxinitrite (Rhee *et al*., 2005; Dietz *et al*, 2006).

Plants respond to attacks by pathogens with rapid increases in reactive oxygen species (R0S) such as superoxide and H_2_0_2_ (Apostol *et al*., 1989). Peroxidases produce R0S that could cause oxidative damage to proteins, DNA, and lipids. Many defects in the immune system of mature *Arabidopsis thaliana* plants with reduced expression of two key peroxidase genes, PRX33 or PRX34, were observed (Daudi *et al*., 2012). Silencing the French-bean class III peroxidase (FBP1) in *A. thaliana* impaired the oxidative burst and rendered plants more susceptible to bacterial and fungal pathogens (Bindschedler *et al*., 2006). Proteomic studies of rice roots seven days after inoculation with *H. seropedicae* showed induction of ascorbate peroxidases (Alberton *et al*., 2013), though the genes encoding these enzymes were not affected in our RNA-seq analyses. Moreover, seven days after inoculation, R0S levels in *Herbaspirillum rubrisubalbicans* attached to rice roots had increased suggesting that the bacteria were subject to oxidative stresses (Valdameri *et al*., 2017). In this work, two peroxidase genes were induced by *H. seropedicae* in inoculated roots.

#### Cell wall

A variety of diazotrophic microorganisms such as *H. seropedicae* Z67, *H*> *rubrisubalbicans* and *A. brasilense* produce cell-wall degrading enzymes (Elbeltagy *et al*., 2001; James *et al*, 2002). Interestingly, the *H. seropedicae* SmR1 genome did not reveal genes coding for known cellulases, pectinases or any other cell-wall degrading enzymes (Pedrosa *et al*., 2011). Nevertheless, rice roots inoculated with *H. seropedicae* induced a gene (2.0-fold) encoding a β-D-xylosidase and repressed 2.4-fold a gene coding for a polygalacturonase suggesting re-modeling of plant cell wall. In *A. thaliana*, β-D-xylosidase has been shown to be involved in secondary cell-wall hemi-cellulose metabolism and plant development (Goujon *et al*., 2003), but little is known about the function of this enzyme.

0ther differentially expressed genes involved in cell-wall metabolism code for proteins similar to an expansin11 (induced 2.6-fold). Expansins have a loosening effect on plant cell-walls and function in cell enlargement as well as in diverse developmental processes in which cell-wall modification occurs including elongation. In addition, they promote elongation of root-hairs (Lin *et al*., 2011; ZhiMing *et al*, 2011) and root-hair initiation (Kwasniewski and Szarejko, 2006).

Expansins have been correlated with plant-bacteria interactions. In tobacco, *Bacillus subtilis* G1, a plant growth promoting bacterium, induced the expression of two expansins NtEXP2 and NtEXP6 (Wang *et al*, 2009). Also, inoculation of *Melilotus alba* with *Sinorhizobium meliloti* lead to enhanced MaEXP1 mRNA levels in roots and nodules (Giordano and Hirsch, 2004). Together, these results suggest that inoculation with *H. seropedicae* also led to modification of plant cell wall which may facilitate bacterial colonization of inner tissue by loosening cell wall.

#### Plant immune responses

The plant immune system can sense and respond to pathogen attacks in two manners: the first involves recognition of pathogen/microbe associated molecular patterns (PAMPs) by surface pattern-recognition receptors (PRRs) resulting in pattern-triggered immunity (PTI). Second, resistance proteins (R) that recognise pathogen effectors are expressed leading to effector-triggered immunity (ETI) (Abramovitch *et al*, 2006; Jones and Dangl, 2006; McDowell and Simon, 2008). Among the regulated genes that have functions involved in defence (in dark gray squares in Figure 3), as determined by MapMan, three genes encoding PR-proteins were repressed (L0C_0s10g25870.1, L0C_0s11g07680.1, L0C_0s02g38392.1, 2.5, 3.4 and 2.7-fold respectively) while one transcript that encoded a lipase called EDS1 (enhanced disease susceptibility 1) was induced 2-fold.

The role of EDS1 in defence is well described in *Arabidopsis* and is required for resistance conditioned by R genes which encode proteins that contain nucleotide-binding sites and leucine-rich repeats (NBS-LLR) (Aarts *et al*., 1998; McHale *et al*., 2006; Bartsch *et al*., 2006; Jones and Dangl, 2006; Venugopal *et al*., 2009). Mutant *eds1* seedlings exhibited enhanced susceptibility to the biotrophic oomycete *Peronospora parasifica* (Parker *et al*, 1996). Analysis of *EDS1*> and *PR-gene* expression showed induction of both genes after inoculation with *Pseudomonas syringae* or treatment with salicylic acid (SA) (Falk *et al*, 1999). In addition, in the *Arabidopsis eds1* mutant, the expression of PR-proteins was undetectable. When the *eds1* mutant was treated with SA however, expression of PR was detected. The authors suggest that EDS1 functions upstream of PR1-mRNA accumulation. Among the 3 genes for PR-proteins repressed, one (L0C_0s02g38392.1) is a NBS-LRR disease resistance protein. Therefore in *H. seropedicae*-rice interaction an inverse correlation between PR-protein and EDS1 expression was observed, perhaps suggesting a fine regulation of these defence systems by *H. seropedicae*.

In previous work down-regulation of genes associated with defence was observed in rice plants inoculated with *H. seropedicae* such as a putative probenazole inducible protein (PBZ1, L0C_0s12g36840.1) that was repressed 3.6-fold as shown by RT-qPCR analysis (Brusamarello-Santos *et al*, 2011). RNA-seq data showed that this gene (L0C_0s12g36840.1) was repressed 2.9-fold. Another two genes coding proteins similar to PBZ1 (L0C_0s12g36830.1 and L0C_0s12g36850.1) were also repressed (7.2 and 4.1, respectively). Kawahara *et al*. (2012) using an RNA-seq approach to study the transcriptome of rice inoculated with the blast fungus *M*> *oryzae* observed that the same PBZ1 genes detected in our study were induced 273 and 233-fold upon inoculation with the pathogen. The induction of PBZ1 gene by pathogens has been considered as a molecular marker for rice defence response (Kim *et al*., 2008).

Another gene regulated by *H. seropedicae* that is related to defence codes for a thionin, a small cysteine-rich protein that occurs in a broad range of plant species (Florack and Stiekema W.J., 1994). Thionins are known for their toxicity to plant pathogens and several studies showed that their over-expression is related to increased resistance to diseases (Iwai *et al*., 2002; Choi *et al*, 2008; Muramoto *et al*., 2012; Ji *et al*, 2015).

Brusamarello-Santos *et al*. (2011) showed 5-fold repression of thionin genes from chromosome 6 seven days after inoculation with *H. seropedicae* but these thionins from chromosome 6 were not regulated in roots three days after inoculation with *H. seropedicae*. Since there are 15 thionin genes (according to the Rice Genome Annotation Project RGAP7) sharing high identity in chromosome 6, we mapped the unmapped reads to the rice genome as well as to chromosome 6 separately and observed an average of 480 mapped reads for control libraries and 136 for inoculated libraries mapping on thionin genes, a result that is in accordance with repression patterns observed in rice roots seven days after inoculation with *H. seropedicae* (Brusamarello-Santos *et al*., 2011). Interestingly, we observed a thionin transcript from chromosome 7 (L0C_0s07g24830.1) that was induced 7.2-fold in the presence of *H. seropedicae* three DAI. Time-dependent regulation of thionin was also observed by Ji *et al*. (2015) in rice roots infected with *Meloidogyne graminicola*. In a by Straub and co-workers study only a few defence-related genes were induced in the transcriptome of *Miscanthus sinensis* inoculated with *Herbaspirillum frisingense* helping to explain why this bacterium can effectively invade and colonise plants (Straub *et al*., 2013).

### Signalling

#### Plant Receptor

The plant immune system employ, at the cell surface, receptors to perceive pathogen associated molecular patterns (PAMPs) or damage associated molecular patterns (DAMPs). Theses receptors have an ectodomain potentially involved in ligand binding, a single transmembrane domain and, most of times, a intracellular kinase domain (Couto and Zipfel, 2016).

Amongst the genes with the highest expression differences seen in RNA-seq data was a wall-associated receptor kinase-like (WAK) 22 precursor gene (L0C_0s10g07556.1), repressed 93-fold (30-fold by RT-qPCR) in H *seropedicae* inoculated roots. Cell wall-associated receptor kinases (WAKs) contain an extracellular domain composed of one or more epidermal growth factor repeats (EGF). Animal proteins containing these repeats are known to bind small peptides (Hynes and MacDonald, 2009). In *A. thaliana* WAKs bind to cross-linked pectin cell wall, pathogen- or damage-induced pectin fragments and oligogalacturonides, thus regulating cell expansion or stress response depending on the state of the pectin (Kohorn and Kohorn, 2012; Kohorn, 2015, 2016). Several studies have described the role of WAK genes in rice resistance to pathogens (Li *et al*., 2009; Cayrol *et al*., 2016; Harkenrider *et al*, 2016; Delteil *et al*, 2016). Plant proteins show a similar domain architecture consisting of cell-wall pectin binding extracellular region, the EGF-like domain, and a kinase domain. Analyses of rice loss-of-function mutants of WAK genes showed that individual genes are important for resistance against *M. oryzae*. 0sWAK14, 0sWAK91 and 0sWAK92 positively regulate resistance while 0sWAK112d is a negative regulator of blast resistance (Delteil *et al*., 2016). Cayrol et al. (2016) demonstrated that 0sWAK14, 0sWAK91 and 0sWAK92 can form homo- and hetero-complexes and hypothesized that the loss of function of any of these proteins may destabilized the complex and affect their functioning. In *A. thaliana* the WAK22 orthologous gene (AtWAKL10) encodes a functional guanylyl cyclase which is co-expressed with pathogen defence related genes (Meier *et al*., 2010). Thus the wall-associated receptor kinase-like 22 gene of rice could be a candidate for surface pattern recognition receptor and its repression may allow *H. seropedicae* to evade activation of the plant-defence system.

A cysteine-rich receptor-like protein kinase (CRK) gene (L0C_0s04g56430.1) and a serine/threonine-protein kinase At1g18390 precursor (L0C_0s05g47770.1) were induced 3.2-fold and fold, respectively, while a lectin-like receptor kinase 7 (L0C_0s07g03790.1) and SHR5-receptor-like kinase (L0C_0s08g10320.1) were repressed 2.7 and 2.5-fold respectively in the presence of *H. seropedicae* (Table S2). The latter protein has 75% identity with sugarcane SHR5 receptor kinase repressed by colonisation with diazotrophic endophytes (Vinagre *et al*., 2006). The authors suggested that the expression levels of this gene were inversely related to the efficiency of beneficial plant-bacterial interactions (Vinagre *et al*, 2006). Besides in *Arabidopsis* cysteine rich receptor-like kinase 5 protein is involved in regulation of growth, development, and acclimatory responses (Burdiak *et al*., 2015).

#### Auxin signalling

Auxins regulate diverse physiological processes such as vascular tissue differentiation, lateral root initiation and have also been linked to defence in plant-pathogen interactions (Rogg and Bartel, 2001; Kazan and Manners, 2009; McSteen, 2010; Adamowski and Friml, 2015). Auxin response elements (AuxREs), when bound to auxin response factors (ARFs), control auxin-dependent gene expression. The Aux/IAA protein family members that inhibit ARFs mediate this regulation (Guilfoyle, 2015).

Four differentially expressed genes related to auxin signalling were identified in rice roots inoculated with *H. seropedicae*. The gene L0C_0s08g24790.1 encoding an auxin-responsive protein was repressed 2.1-fold (Table 2) whereas the genes coding for auxin-induced proteins, L0C_0s09g25770.1 and L0C_0s05g01570.1, were repressed 1.8 and 2.5-fold, respectively. Repression of genes related to auxin signalling was reported in rice roots inoculated with *H. seropedicae* (Brusamarello-Santos et al. 2011). Brusamarello-Santos et al. (2011) found that ARF2-like, IAA 11 and IAA18 were repressed, 1.4, 1.5 and 2.8-fold respectively, in the presence of *H. seropedicae* 7 days after inoculation. These genes were not regulated in our data probably due to the time difference of the cDNA library construction and too low expression levels.

**Table 2.**
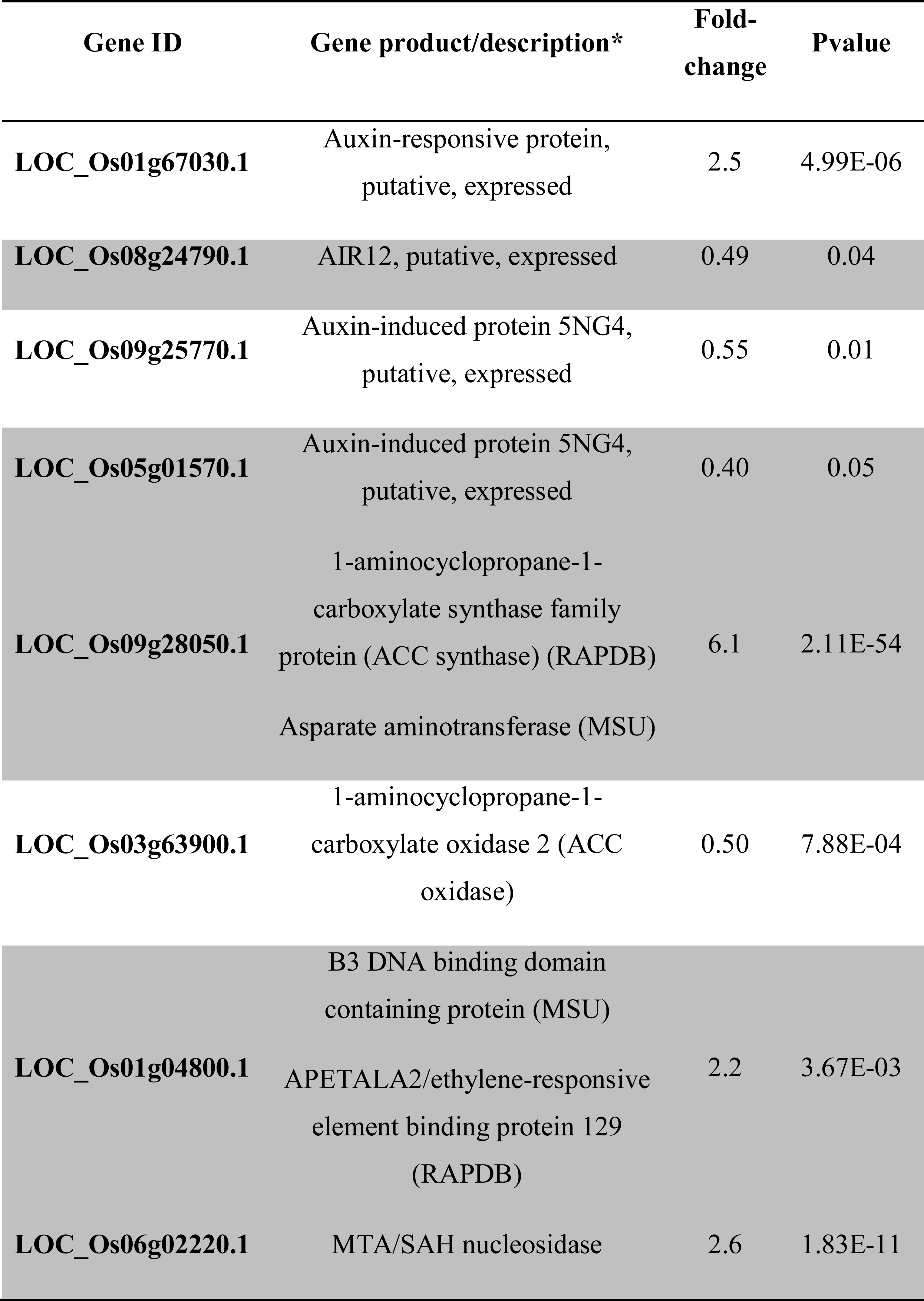
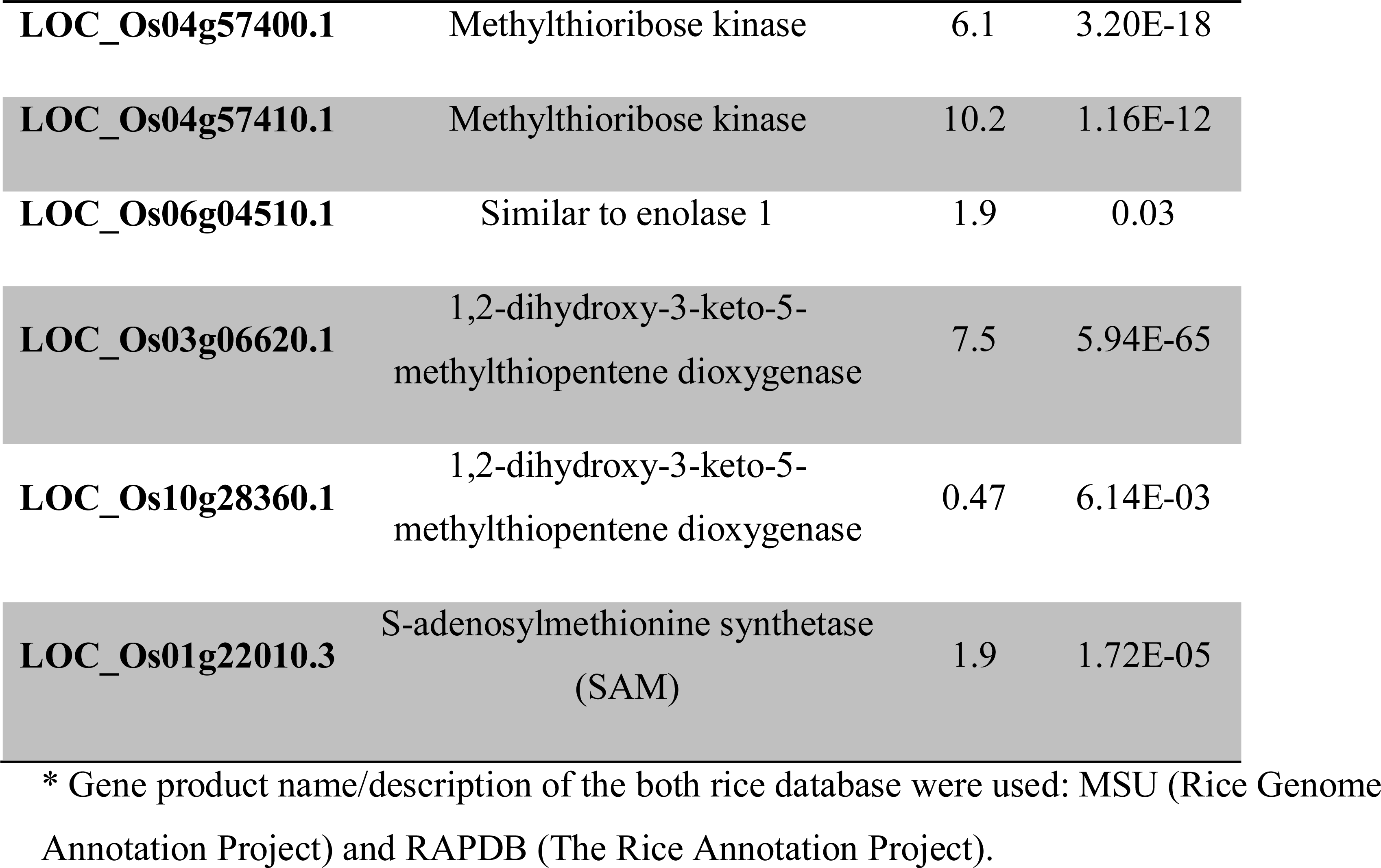
Rice genes modulated by colonisation with *H. seropedicae* that are involved in phyto-hormone signalling and the MTA cycle

*H. seropedicae* can produce IAA in the presence of tryptophan, thus suggesting the plant growth promotion effect may be due to bacterial derived auxin (Bastian *et al*., 1998). In addition exogenous application of auxin lead to increase in lateral root numbers (Inukai *et al*., 2005). However the number of lateral roots in rice plants inoculated and non-inoculated with *H. seropedicae*, was determined (data not shown) but no significant difference was observed.

In the absence of clear regulation of auxin-dependent genes and phenotype, the results suggest that auxin does not play an important role in Nipponbare rice roots colonization by *H. seropedicae*. 0n the other hand the repression of auxin signalling has been correlated to defence responses. Auxin levels have been correlated with susceptibility to pathogens (Kazan and Manners, 2009) that are able to produce high levels of auxin (Jameson, 2000). It has also been shown that pathogen-associated molecular patterns (PAMPs) induce the expression of a miRNA that negatively regulates mRNAs for F-box auxin receptors leading to resistance to *P. syringae* in *Arabidopsis* (Navarro *et al*, 2006). The observed repression in several genes involved with defence in the rice-*H*> *seropedicae* interaction opens the question whether auxin could be important for bacteria survival inside the plant by down-regulation defence system. Further studies are needed to elucidate if and how auxin signalling participates in plant-bacterial interactions.

#### Ethylene signalling

Ethylene is also involved in several biological processes that activate defence responses and adventitious root-growth in rice and other plants (Lorbiecke and Sauter, 1999; Glazebrook, 2005; Robert-Seilaniantz *et al*., 2011). Ethylene can be synthesised by oxidation of 1-aminocyclopropane-1-carboxylate (ACC) by ACC oxidase. ACC is synthesised from adenosylmethionine (AdoMet) by ACC synthase (ACS). ACS is up regulated 6-fold and ACC oxidase is repressed 2-fold by inoculation with *H. seropedicae*. An ethylene response factor (ERF) (L0C_0s01g04800.1) gene was also induced 2.2- fold in inoculated roots. These data indicate that ethylene synthesis is attenuated in the presence of bacteria. Moreover Alberton et al. (2013) measured the level of ethylene in inoculated rice roots (seven days after inoculation) and found a decrease of ethylene levels. Furthermore, Valdameri *et al*. (2017) detected induction of ACC oxidase in rice plants inoculated with the pathogen H *rubrisubalbicans*. These results suggest that the ethylene pathway is differentially modulated in the presence of pathogens and beneficial endophytic bacteria that promote plant growth.

#### SA signalling

Salicylic acid is derived from phenolic compounds and is involved in response to attack by pathogens (Vlot *et al*, 2009; An and Mou, 2011). In rice roots inoculated with *H. seropedicae*, a SA-dependent carboxyl methyltransferase family protein (L0C_0s11g15340.2) was repressed 40-fold. A member of this family is salicylic acid carboxyl methyltransferase (SAMT) that catalyses the formation of methyl salicylate (MeSA) from SA (Ross *et al*., 1999). MeSA is an essential signal for systemic acquired resistance (SAR) in tobacco plants. In addition mutations in SAMT showed that this gene is required for SAR (Park *et al*., 2007).

SA signalling is differentially regulated by members of the WRKY transcription factor family (Eulgem and Somssich, 2007). In inoculated rice roots, two WRKY transcription factors were regulated, one repressed (2.3-fold) and one induced (2.0-fold). The induced gene encodes a WRKY51 similar to WRKY11 of *Arabidopsis*, whereas the repressed gene encodes WRKY46, which was shown to be induced by SA in *Arabidopsis*. WRKY11 is a negative regulator of resistance (Journot-Catalino *et al*, 2006) and *Arabidopsis* plants in which WRKY46 was over-expressed were more resistant to *P. syringae* (Hu *et al*., 2012). These results are in agreement with the hypothesis that the SA signalling and defence system are attenuated in the presence of the *H. seropedicae*.

### Metal ion metabolism

Several genes related to metal transport were differentially expressed, most of them up regulated in the presence of *H. seropedicae*. Amongst the 20 most highly regulated rice genes, eight were related to metal transport (Table 3; Table 4). Phyto-siderophore synthesis starts with production of nicotianamine (NA) from S-adenosylmethionine, which in turn is derived from 5’-methylthioadenosine of the methionine salvage pathway (MTA cycle) (Figure 6). Transcripts of enzymes of the MTA cycle encoding SAM, MTA nucleosidase, MTR kinase, E1 enolase/phosphatase and acireductone dioxygenase (ARD) were induced 1.9, 2.6, 6.0, 1.9 and 7.5-times, respectively, in roots colonised by *H*> *seropedicae* (Figure 6). Using proteomic and RT-qPCR analyses, Alberton (2013) observed similar expression patterns in rice roots inoculated with *H. seropedicae*.

**Figure 6:**
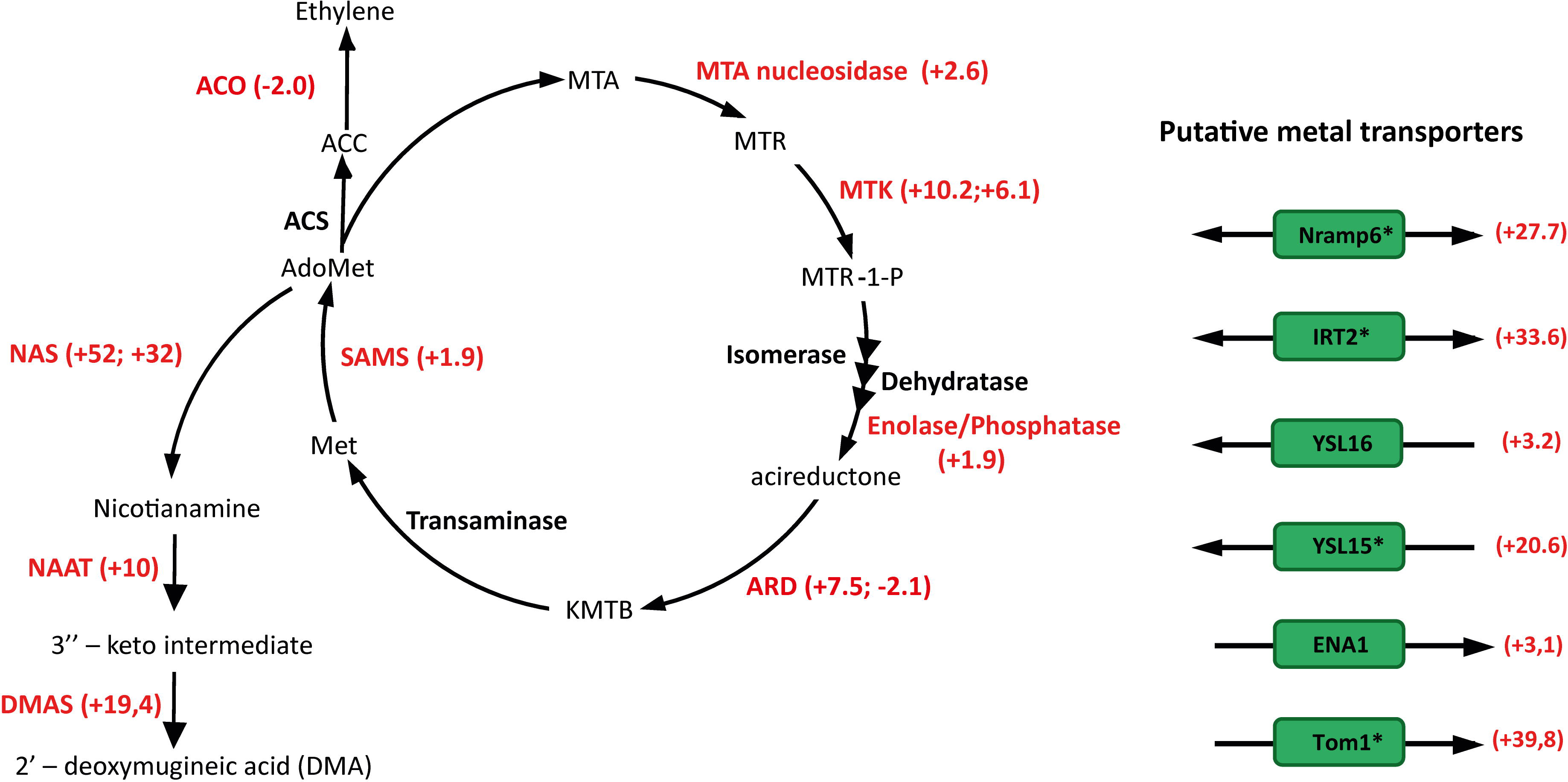
Differentially expressed genes in rice roots following colonisation by *H. seropedicae*. Genes involved in siderophore synthesis and transport, the methionine salvage pathway and ethylene synthesis are shown. Numbers in parentheses represent the fold change. *H. seropedicae* SmR1 induces methionine recycling and mugineic acid (MA) synthesis as well as the expression of transporters involved in iron metabolism. The expression of those genes marked with an asterisk was confirmed by RT-qPCR Abbreviations: AdoMet, S-adenosylmethionine; ACC, 1-aminocyclopropane-1-carboxylate; ACS, 1-aminocyclopropane-1-carboxylate synthase; ACO, 1-aminocyclopropane-1-carboxylate oxidase; MTA, 5'-methylthioadenosine; MTR, 5'-methylthioribose; MTK, methylthioribose kinase; MTR-1-P, 5'-methylthioribose-1-phosphate; KMTB, 2-keto-4-methylthiobutyrate; ARD, acireductone dioxygenase; SAMS, S-adenosylmethionine synthetase; NAS, nicotianamine synthase; NAAT nicotianamine aminotransferase; DMAS, deoxymugineic acid synthase; Tom1, transporter of mugineic acid 1; ENA1 (efflux transporters of nicotianamine 1); Nramp6, Natural Resistance-Associated Macrophage Protein; IRT2(iron-regulated transporter 2); YSL16 (yellow strip-like gene 16); YSL15 (yellow strip-like gene 15)

**Table 3.**
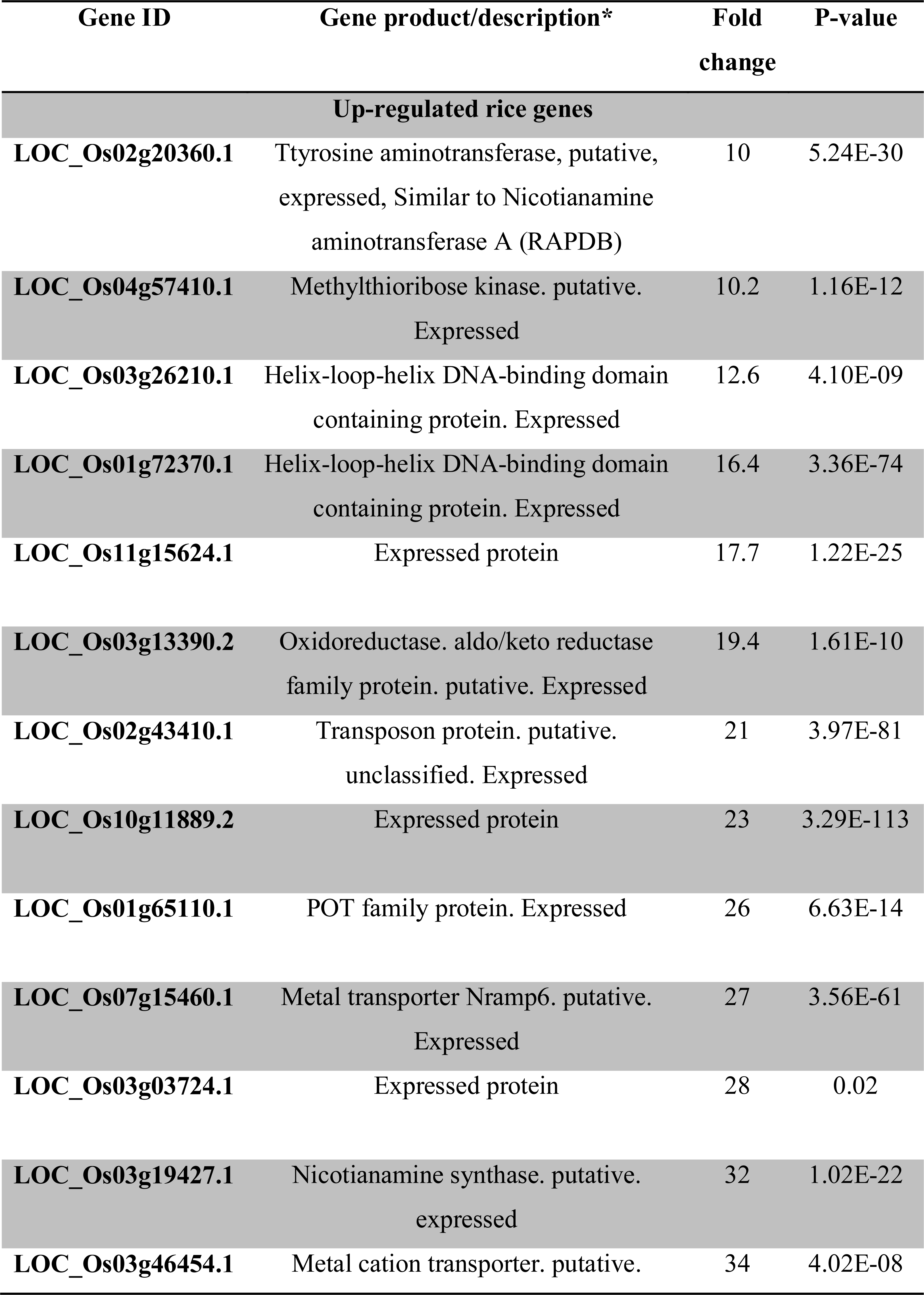
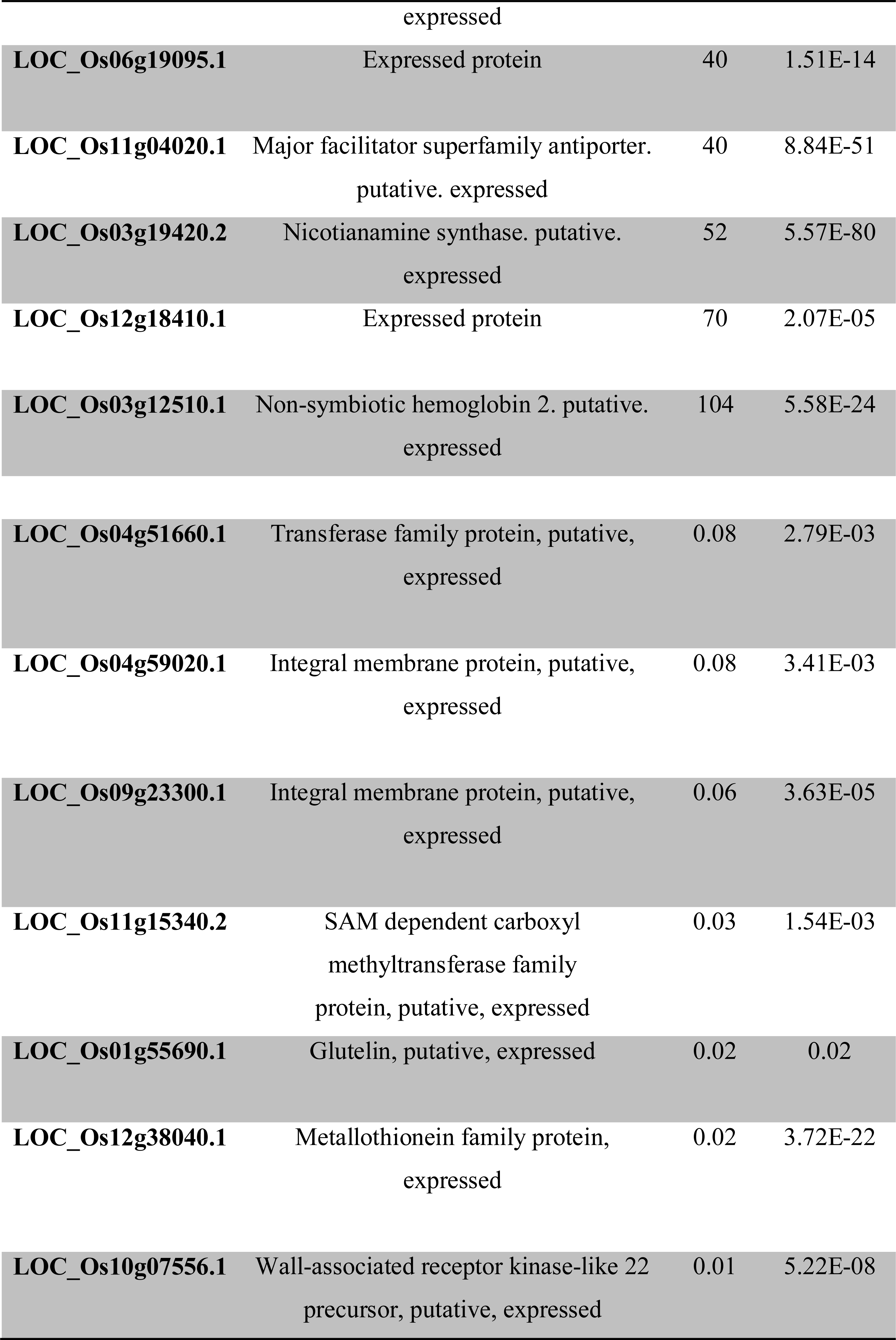
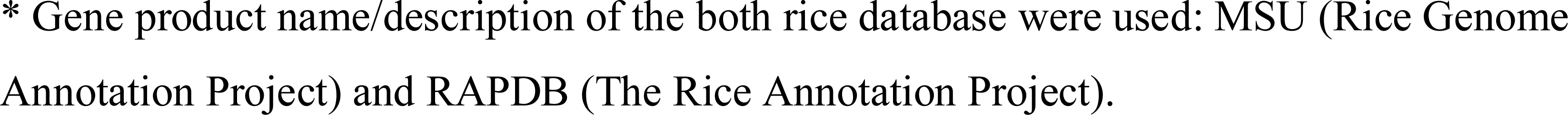
Rice genes highly regulated (fold change > 10) by inoculation with *H. seropedicae*

**Table 4:**
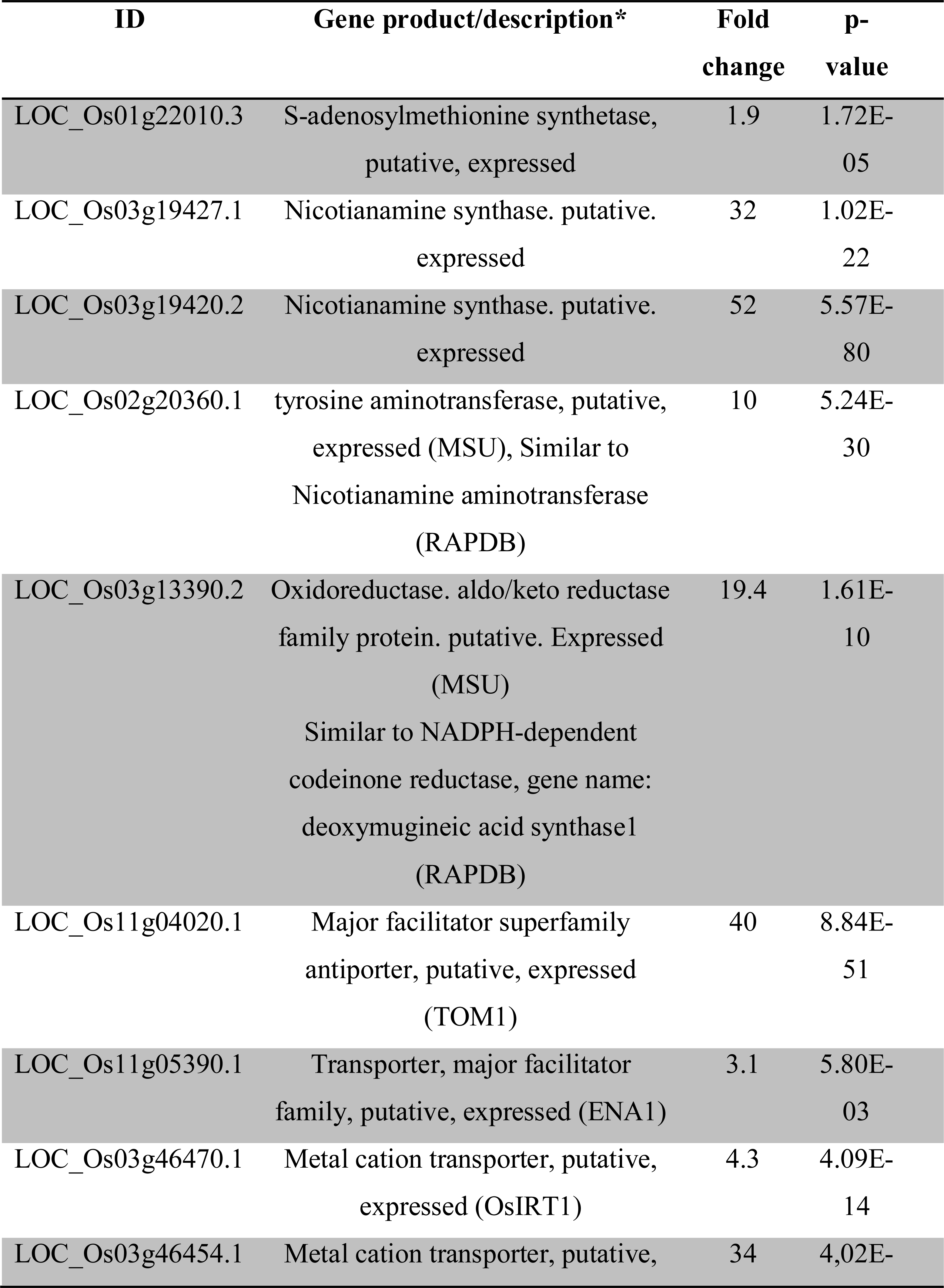
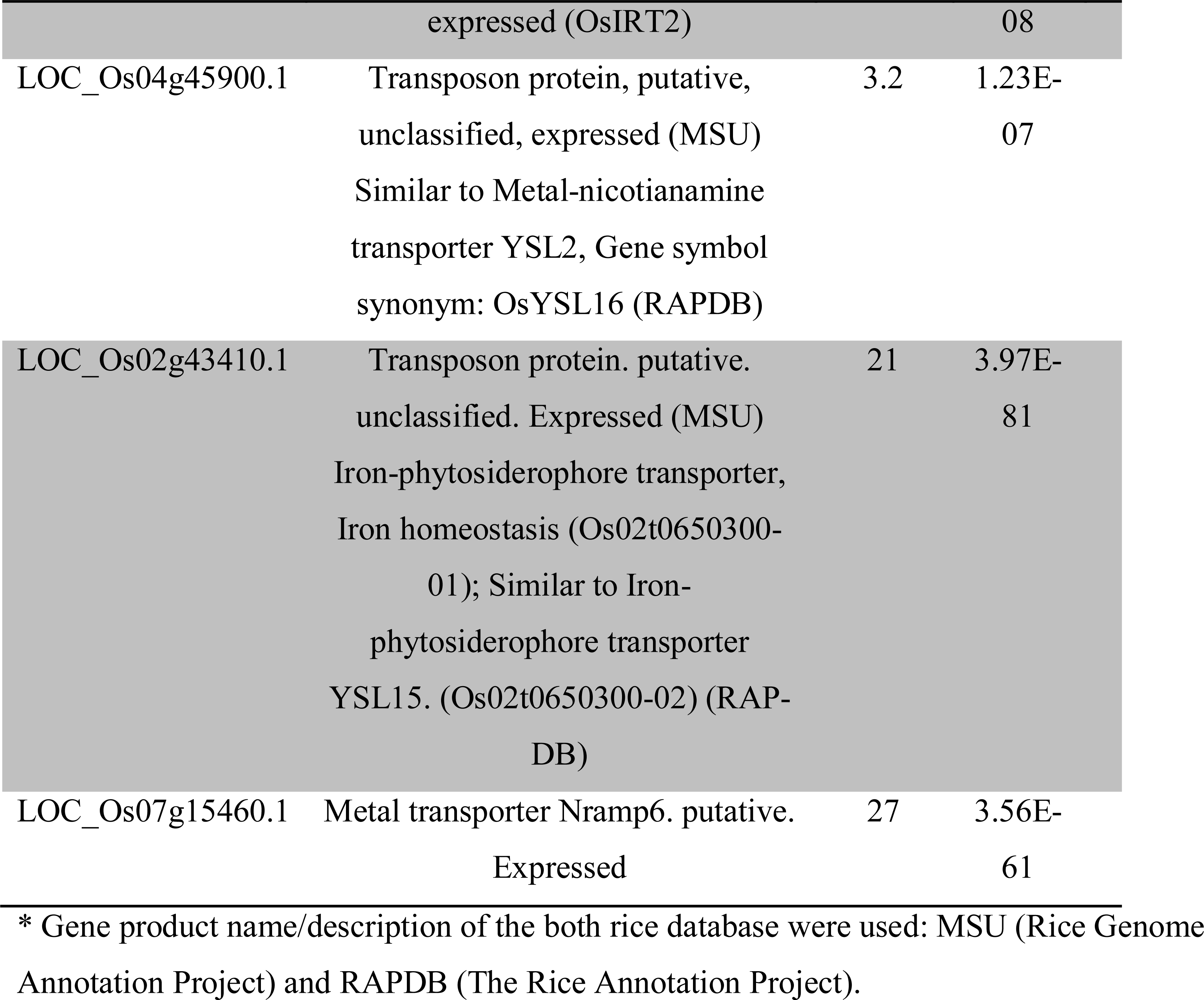
Differentially expressed genes in rice roots colonised by *H. seropedicae* involved in uptake and transport of metals

| **ID** | **Gene product/description\*** | **Fold change** | **p-value** |
| --- | --- | --- | --- |
| LOC_Os01g22010.3 | S-adenosylmethionine synthetase, putative, expressed | 1.9 | 1.72E-05 |
| LOC_Os03g19427.1 | Nicotianamine synthase. putative. expressed | 32 | 1.02E-22 |
| LOC_Os03g19420.2 | Nicotianamine synthase. putative. expressed | 52 | 5.57E-80 |
| LOC_Os02g20360.1 | tyrosine aminotransferase, putative, expressed (MSU), Similar to Nicotianamine aminotransferase (RAPDB) | 10 | 5.24E-30 |
| LOC_Os03g13390.2 | Oxidoreductase. aldo/keto reductase family protein. putative. Expressed (MSU)<br>Similar to NADPH-dependent codeinone reductase, gene name: deoxymugineic acid synthase1 (RAPDB) | 19.4 | 1.61E-10 |
| LOC_Os11g04020.1 | Major facilitator superfamily antiporter, putative, expressed (TOM1) | 40 | 8.84E-51 |
| LOC_Os11g05390.1 | Transporter, major facilitator family, putative, expressed (ENA1) | 3.1 | 5.80E-03 |
| LOC_Os03g46470.1 | Metal cation transporter, putative, expressed (OsIRT1) | 4.3 | 4.09E-14 |
| LOC_Os03g46454.1 | Metal cation transporter, putative, | 34 | 4,02E- |
|  | expressed (OsIRT2) |  | 08 |
| LOC_Os04g45900.1 | Transposon protein, putative,<br>unclassified, expressed (MSU)<br>Similar to Metal-nicotianamine<br>transporter YSL2, Gene symbol<br>synonym: OsYSL16 (RAPDB) | 3.2 | 1.23E-07 |
| LOC_Os02g43410.1 | Transposon protein. putative.<br>unclassified. Expressed (MSU)<br>Iron-phytosiderophore transporter,<br>Iron homeostasis (Os02t0650300-01); Similar to Iron-phytosiderophore transporter<br>YSL15. (Os02t0650300-02) (RAP-DB) | 21 | 3.97E-81 |
| LOC_Os07g15460.1 | Metal transporter Nramp6. putative.<br>Expressed | 27 | 3.56E-61 |

\* Gene product name/description of the both rice database were used: MSU (Rice Genome Annotation Project) and RAPDB (The Rice Annotation Project).

The synthesis of NA and MAs involves a set of enzymes including S-adenosylmethionine syntase (SAM) that catalyses the adenylation of L-methionine to S-adenosylmethionine, nicotianamine synthase (NAS) that converts S-adenosylmethionine to nicotianamine, and nicotianamine aminotransferase (NAAT) that catalyses the amino transfer of NA to produce the 3’’-keto intermediate that is reduced by deoxymugineic acid synthase (DMAS) to produce 2’-deoxymugineic acid (DMA) (Shojima *et al*., 1990; Higuchi *et al*., 1999; Bashir *et al*., 2006; Inoue *et al*., 2008).

Two nicotianamine synthase (NAS) genes of rice - L0C_0s03g19420.2 and L0C_0s03g19427.1 were induced (52- and 32-fold, respectively) in inoculated rice roots. L0C_0s03g19427.1 was also induced in rice roots inoculated with *Azospirillum spp*. (Drogue *et al*., 2014). Increased levels of nicotianamine have been shown to increase Fe uptake in rice plants (Lee *et al*, 2009). Furthermore, in *Lotus japonicus* inoculated with *Mesorhizobium loti*, nicotianamine synthase 2 was expressed only in nodules pointing to a role in symbiotic nitrogen fixation (Hakoyama *et al*., 2009). A tomato mutant defective in the synthesis of nicotianamine was affected in iron metabolism (Ling *et al*., 1996). In addition NAAT (L0C_0s02g20360.1) and DMAS (L0C_0s03g13390.2) were also induced 10-fold and 19-times respectively in colonised rice roots. Iron deficiency provoked induction of rice gene 0sNAAT1 (Inoue *et al*., 2008).

Recently, members of a major facilitator super-family have been described as essential to the efflux of MA and NA in rice. T0M1 (transporter of mugineic acid 1) is involved in the efflux of MA, while ENA1 (efflux transporters of nicotianamine 1) and ENA2 in the efflux of NA (Nozoye *et al*, 2011). T0M1 (L0C_0s 11 g04020.1) and ENA1 (L0C_0s11g05390.1) were induced 40 and 3-fold in colonised roots. Induction of T0M1 was confirmed by RT-qPCR (Figure 4).

Fe^+++^ ions chelated by PS need to be transported inside the cell. In gramineous plants, two groups of Fe-MA transporters are present: ZmYS1 (Curie *et al*., 2001) and the YSL (yellow strip-like) transporter family (Inoue *et al*., 2009). Inoue et al. (2009) analysed the expression of 18 YSL genes in rice and observed induction of 0sYSL15 and 0SYSL16 genes under iron-deficiency. 0ther studies have shown that 0sYSL2 (L0C_0s02g43370) takes up Fe^2+^-NA (Ishimaru *et al*., 2010) and 0sYSL16 Fe^3+^-DMA (L0C_0s04g45900.1) (Kakei *et al*, 2012). Here we showed induction (3.2-fold) of 0sYSL16 (L0C_0s04g45900.1) and 21-fold increase of transcripts encoding a gene similar to 0SYSL15 (L0C_0s02g43410.1). Induction of 0SYSL15 was confirmed by RT-qPCR (Figure 4). Furthermore, 0sIRT1 (iron-regulated transporter 1) and 0sIRT2 were also induced under low Fe conditions (Ishimaru *et al*., 2006). We detected induction of 0sIRT2 (L0C_0s03g46454.1) (34-fold in RNA-seq and 93-fold in RT-qPCR) (Figure4).

Other metal transporters such as Nramp6 (L0C_0s07g15460.1) were also up regulated (27fold) in colonised rice roots, a result confirmed by RT-qPCR (Figure 4). Members of the natural resistance-associated macrophage protein (NRAMP) family are transition metal cation/proton cotransporters or anti-porters of broad specificity. AtNRAMP6 of *Arabidopsis* is up-regulated in response to iron deficiency and is involved with metal mobilization from vacuoles to cytosol (Thomine *et al*, 2003). Induction of Nramp6 in rice roots colonised by *H. seropedicae*. In a previous study in rice inoculated with *Azospirillum* spp the expression of this gene was also induced (Drogue et al. 2014). In addition a gene for an integral membrane protein (L0C_0s09g23300.1) named 0sVIT2 was 17.7-fold repressed by *H. seropedicae*. This gene is involved in transport of Fe/Zn into vacuoles and is up-regulated in rice roots with excess Fe. Knockout/ knockdown of this gene led to Fe accumulation in seeds (Zhang *et al*., 2012; Bashir *et al*., 2013). Together, these results suggest that colonised roots respond in such a manner as to accumulate Fe.

Interestingly the transcript with the highest fold change [104-times - a result confirmed using RT-qPCR (Figure 4)] in colonised roots codes for the non-symbiotic hemoglobin 2 (L0C_0s03g12510.1). High levels of non-symbiotic hemoglobin 2 could help buffer free oxygen and protect bacterial nitrogenase. Arredondo-Peter *et al*. (1997) described two hemoglobins, Hb1 and Hb2, in rice. The Hb2 induced by *H. seropedicae* is very similar to the one described by Arredondo-Peter et al. (1997) (coverage 82 % with an identity of 97 %). In addition, Lira-Ruan *et al*. (2001) studied the synthesis of hemoglobins in rice under normal and stress conditions, coming to the conclusion that Hbs are not part of a generalised stress response. They demonstrated that Hb1 is expressed in different rice organs (root and leaves) during plant development. In etiolated rice plants under 02 limiting conditions the Hb levels increase (Lira-Ruan et al. 2001). This increase suggest that Hb expression maybe due to reduced 02 levels in the presence of the bacteria which make the root environment microaerophilic. The higher requirement for Fe needed for incorporation into Hb may partially explain activation of siderophore synthesis and Fe accumulation. We also found a *H. seropedicae* bacterioferritin (Hsero_1195) gene induced (2.2-fold), but symptoms of Fe deficiency in colonised rice plants were not observed.

Iron homeostasis has been related to plant defence, R0S accumulation and immunity. Also, Fe deficiency triggers accumulation of antimicrobial phenolics compounds. It has been suggested recently that Fe sequestration by bacterial siderophore could be a signal for pathogen infection (Aznar *et al*., 2015). However, bacterial genes involved in siderophore biosynthesis were not observed among the *H. seropedicae* genes expressed in rice roots. Also, transcriptomic analysis of *H. seropedicae* attached to wheat and mayze roots did not show iron metabolism genes up-regulated (Pankievicz *et al*., 2016; Balsanelli *et al*., 2016). These results suggest that the effect of bacteria on plant iron mebabolism is more complex than those caused by iron sequestration.

### Transcripts expressed in interactions with bacteria

The libraries from inoculated roots were mapped against the *H*> *seropedicae* genome (24,278 reads). Amongst the 4,085 annotated genes, 287 had at least one-fold coverage (this set of genes was called *H. seropedicae* expressed genes). Functional classification of expressed genes was performed using the C0G system. After ribosomal genes, the most abundant functional classes found were “unknown”, “energy production and conversion”, “amino acid transport and metabolism”, “cell motility and cell wall” (Table 5). Comparison of expressed genes of *H*> *seropedicae* detected in plants with bacterial genes expressed in culture revealed only 16 differences [p-value < 0.05 using the DESeq statistical package (Table 5)].

**Table 5.** 0RFs of *H. seropedicae* expressed following contact with rice roots

| ID | Fold Change | p-value | ID Feature | Description | COG |
| --- | --- | --- | --- | --- | --- |
| **Hsero_3499** | 0.4 | 7.0E-04 | qor | qor<br>NADPH:quinone<br>oxidoreductase<br>protein<br>4007887:4008915<br>forward | C;R - Energy<br>production and<br>conversion;General<br>function prediction<br>only |
| **Hsero_1gy130** | 30 | 6.20E-03 | Hsero_1130 | Hsero_1130 ABC-type dipeptide<br>transporter,<br>periplasmic<br>peptide-binding<br>protein<br>1287409:1289031<br>forward | E - Amino acid<br>transport and<br>metabolism |
| **Hsero_4713** | 27 | 7.70E-03 | urtA | urtA ABC-type urea<br>transport system,<br>periplasmic<br>component protein<br>5407999:5409252<br>forward | E - Amino acid<br>transport and<br>metabolism |
| **Hsero_0084** | 43 | 0.02 | glnK | glnK nitrogen<br>regulatory PII-like<br>protein<br>99052:99390<br>forward | E - Amino acid<br>transport and<br>metabolism |
| **Hsero_1469** | 0.2 | 2.0E-04 | ttuC | ttuC tartrate dehydrogenase protein<br>1690575:1691654<br>reverse | G - Carbohydrate transport and metabolism |
| **Hsero_1123** | 84 | 0.02 | Hsero_1123 | Hsero_1123 family<br>II aminotransferase protein<br>1278563:1279939<br>reverse | H - Coenzyme transport and metabolism |
| **Hsero_1509** | 0.1 | 2.0E-03 | rplS | rplS 50S ribosomal subunit protein L19<br>1731532:1731915<br>forward | J - Translation, ribosomal structure and biogenesis |
| **Hsero_0677** | 0.3 | 0.02 | ompW2 | ompW2 outer membrane W protein<br>741190:741927<br>forward | M - Cell wall |
| **Hsero_2580** | 0.4 | 5.0E-03 | lon | lon ATP-dependent protease LA protein<br>2940893:2943301<br>reverse | O - Posttranslational modification, protein turnover, chaperones |
| **Hsero_0085** | 20 | 0.04 | amtB | amtB ammonium transporter transmembrane protein<br>99406:100938<br>forward | P - Inorganic ion transport and metabolism |
| **Hsero_2905** | 0.1 | 1.10E-03 | Hsero_2905 | Hsero_2905<br>conserved<br>hypothetical<br>protein<br>3301039:3301272<br>reverse | S - Function<br>unknown |
| **Hsero_0083** | 15 | 0.03 | Hsero_0083 | Hsero_0083<br>membrane protein<br>98251:99039<br>forward | S - Function<br>unknown |
| **Hsero_2723** | 90 | 1.90E-03 | Hsero_2723 | Hsero_2723 methyl-accepting<br>chemotaxis<br>transmembrane<br>protein<br>3101951:3103597<br>reverse | T;N - Signal<br>transduction<br>mechanisms;Cell<br>motility |
| **Hsero_1215** | 0.3 | 0.04 | trnK | trnK tRNA-Lys<br>1374570:1374645<br>reverse | trnK |
| **Hsero_1466** | 0.2 | 1.7E-03 | trnL | trnL tRNA-Leu<br>1689660:1689744<br>reverse | trnL |
| **Hsero_1546** | 0.1 | 0.03 | trnS | trnS tRNA-Ser<br>1781783:1781873<br>forward | trnS |

Genes involved in nitrogen fixation *(nif)* were found amongst genes classified as “energy production and conversion” and “amino acid transport and metabolism”. *Nif-genes* encode proteins involved in the synthesis, maturation and assembly of the nitrogenase complex (Chubatsu *et al*., 2012). *nifD* and *nifH*(coverage 1.7 and 1.2 respectively) were highly expressed in *H*> *seropedicae* colonising wheat and maize roots (Pankievicz *et al*, 2016; Balsanelli *et al*, 2016). We also found two ferredoxin genes important for nitrogenase activity: *fdxA* and *fdxN*(coverage 1.9 and 1.6) (Souza *et al*., 2010).

The promoters of *nif* genes are activated by NifA that is regulated by the Ntr system. When comparisons were made between free-living *H. seropedicae* and those grown in association with rice, *glnK* and *amtB* of the Ntr system were induced 43 and 20-times respectively in the plant-bacterial interaction. Pankievicz *et al*. (2016) also found that *amtB* was induced in *H*> *seropedicae* attached to maize roots. Furthermore, *fixN*and *fixP* (both 1.6X coverage) were also detected *in planta* along with an *urtA* ABC-type urea transport system (Hsero_4713) (27-fold of induction *in planta)*.

Twenty-three genes related to cell motility were found, four with > 2X coverage. Amongst these were *cheW*(a positive regulator of CheA), *ñhD, fliC*and *pilZ*(Hsero_2062) that encodes a type pilus assembly protein. A methyl-accepting chemotaxis trans-membrane protein (Hsero_2723) was induced 90-fold when compared with expression in culture (Tadra-Sfeir *et al*., 2015). This gene was also found to be up-regulated in epiphytic *H seropedicae* colonising wheat and maize (Pankievicz *et al*., 2016; Balsanelli *et al*., 2016). Methyl-accepting chemotaxis proteins interact with Che proteins to detect signals from the environment. Although other T4SS genes were not found perhaps these data suggest that type IV secretion plays a role in *H-seropedicae-rice* interactions. In the nitrogen-fixing, endophytic bacterium *Azoarcus* spp., mutation of type IV pilus *pilAB* genes negatively affects colonisation of rice roots (Dörr *et al*, 1998).

Among the 19 cell-wall related genes, seven were covered at least 2X and an outer-membrane porin (Hsero_4295) was induced 28X. Another membrane-protein (Hsero_0083) of unknown function was induced 14-fold. These could be proteins that *H seropedicae* uses to recognise rice.

## Supplementary Data

Table S1 - 0ligonucleotides used in this research

Table S2 - Genes involved in secondary metabolism and abiotic and biotic stresses modulated in rice roots colonised with *H. seropedicae*.

Table S3 - Differentially expressed genes in rice roots colonised by *H. seropedicae*

## Acknowledgments

We are grateful M.G. Yates for critical reading of the manuscript, Leonardo M. Cruz and Rodrigo A. Cardoso for bioinformatics support. and Instituto Riograndense do Arroz (IRGA) for providing seeds. We are also thankful to Roseli Prado, Marilza Doroti Lamour and Valter A. Baura for technical assistance. This work was supported by National Institute of Science and Technology of Nitrogen Fixation (INCT - Nitrogen Fixation), Council for Scientific and Technological Development (CNPq) and Coordination of Improvement of Higher-Education Personnel (CAPES). Thanks to CNPq for doctoral scholarship.

